# Polar localization of putative phospholipid transporters in *Escherichia coli*

**DOI:** 10.1101/2025.04.17.649288

**Authors:** Wee Boon Tan, Zhi-Soon Chong, Jacob Wye Meng Cheong, Jiang Yeow, Shu-Sin Chng

**Affiliations:** Department of Chemistry, National University of Singapore, Singapore 117543; Singapore Center for Environmental Life Sciences Engineering, National University of Singapore (SCELSE-NUS), Singapore 117456

**Author notes:** These authors contributed equally (Wee Boon Tan, Zhi-Soon Chong, Jacob Wye Meng Cheong). **Author Contributions:** W.B.T., Z.-S.C, J.W.M.C, and J.Y. performed the experiments and initial data analysis. W.B.T. and S.-S.C. wrote the manuscript. **Competing Interest Statement:** The authors declare no competing interests.

**Keywords:** Outer membrane, YhdP, TamB, YdbH, bridge-like lipid transfer protein

## Abstract

The Gram-negative bacterial cell envelope comprises an outer membrane (OM) with an asymmetric arrangement of lipopolysaccharides (LPS) and phospholipids (PLs), protecting them from both physical and chemical threats. To build the OM, PLs must be transported across the cell envelope; this process has remained elusive until recently, where three collectively essential AsmA-superfamily proteins – YhdP, TamB, and YdbH – are proposed to function as anterograde PL transporters in Escherichia coli. Here, we identify the cell wall-binding protein DedD as a novel interacting partner of YhdP, and discover that all three AsmA-superfamily proteins are recruited to, and strongly enriched at the cell poles. Our observation raises the possibility that anterograde PL transport could be spatially restricted to the cell poles, and highlights the importance of understanding the spatial-temporal regulation of OM biogenesis in coordination with cell growth and division.

## Importance

The outer membrane of Gram-negative bacteria serves as an effective permeability barrier, and confers intrinsic antibiotic resistance. This barrier function requires distinct distribution of lipids across the bilayer, yet how phospholipids, the most basic building block, get transported and assembled into the outer membrane is not well understood. In this study, we describe the observation revealing that three putative phospholipid transporters are mostly present at the cell poles in *Escherichia coli*, highlighting possible polar localization of lipid transport to ultimately support outer membrane biogenesis during growth and division. Our work sets the stage for studying how phospholipid transport impacts OM stability, lipid asymmetry, and/or function, thus informing future strategies for antibiotics development against these processes.

## Observation

The outer membrane (OM) is an essential barrier that protects Gram-negative bacteria from antibiotics and detergents (1). Critical to this function is the asymmetric arrangement of lipids in the OM, which is presumably established via the direct placement of lipopolysaccharides (LPS) and phospholipids (PLs) into the outer and inner leaflets, respectively (2). LPS is transported from the inner membrane (IM) to the OM along the continuous hydrophobic groove of the seven-component Lpt bridge, eventually inserted into the outer leaflet all around the cells except the poles (3, 4). By comparison, how PLs are moved bidirectionally across the cell envelope has been less clear (2). Recently, it was proposed that three AsmA-superfamily proteins, YhdP, TamB, and YdbH, are involved in this essential process of anterograde PL transport in *Escherichia coli* (5, 6). These trans-envelope proteins are structural homologs of eukaryotic bridge-like lipid transport proteins, featuring repeating β-groove motifs along their predicted/solved structures compatible with lipid binding and transport (7, 8). Indeed, experimental and *in silico* evidences that support PL binding in these hydrophobic grooves are emerging (9–11), though mechanistic understanding of transport is lacking. For example, it is not clear what other factor(s) might be required as part of the functional complexes of these AsmA-superfamily proteins, nor is it known where these complexes may assemble in the cell. YdbH interacts with the OM lipoprotein YnbE (12), and TamB interacts with the OM β-barrel protein (OMP) TamA, the latter also in fact has an established function in OMP assembly (13, 14). In this regard, much less is known about YhdP.

### YhdP interacts with the cell division protein DedD

To identify potential interacting partner(s) of YhdP, we expressed a functional C-terminally 8xHis-tagged version of YhdP at low levels from a pET23/42 vector in an *E. coli* wild-type strain (Figure S1), and performed affinity purification after isolating and solubilizing the total membrane fraction. The pET23/42 plasmid allows “leaky” expression of proteins in the absence of the T7 RNA polymerase (15), and has previously been used to mimic native protein expression levels to study cell envelope biology (12, 15–17). YhdP-His migrated as a ~150 kDa band on SDS-PAGE (Figure 1A). An additional band between 35-40 kDa was readily observed only in the sample containing YhdP-His (Figure 1A), suggesting a possible interacting partner. Interestingly, tandem mass spectrometry identified this co-purified band as the cell division protein DedD. We could no longer detect this unique band in the Δ*dedD* strain (Figure 1A). To further validate this interaction, we performed reciprocal purification, and demonstrated that N-terminally 6xHis-tagged DedD pulled down C-terminally 3xFLAG-tagged YhdP (Figure 1B, S1). We were also able to predict a possible structural model of a YhdP-DedD heterodimer using AlphaFold3 (18), revealing topologically sound interactions at both the transmembrane and periplasmic regions (Figure S2). We conclude that YhdP specifically interacts with DedD in cells.

**Figure 1.**
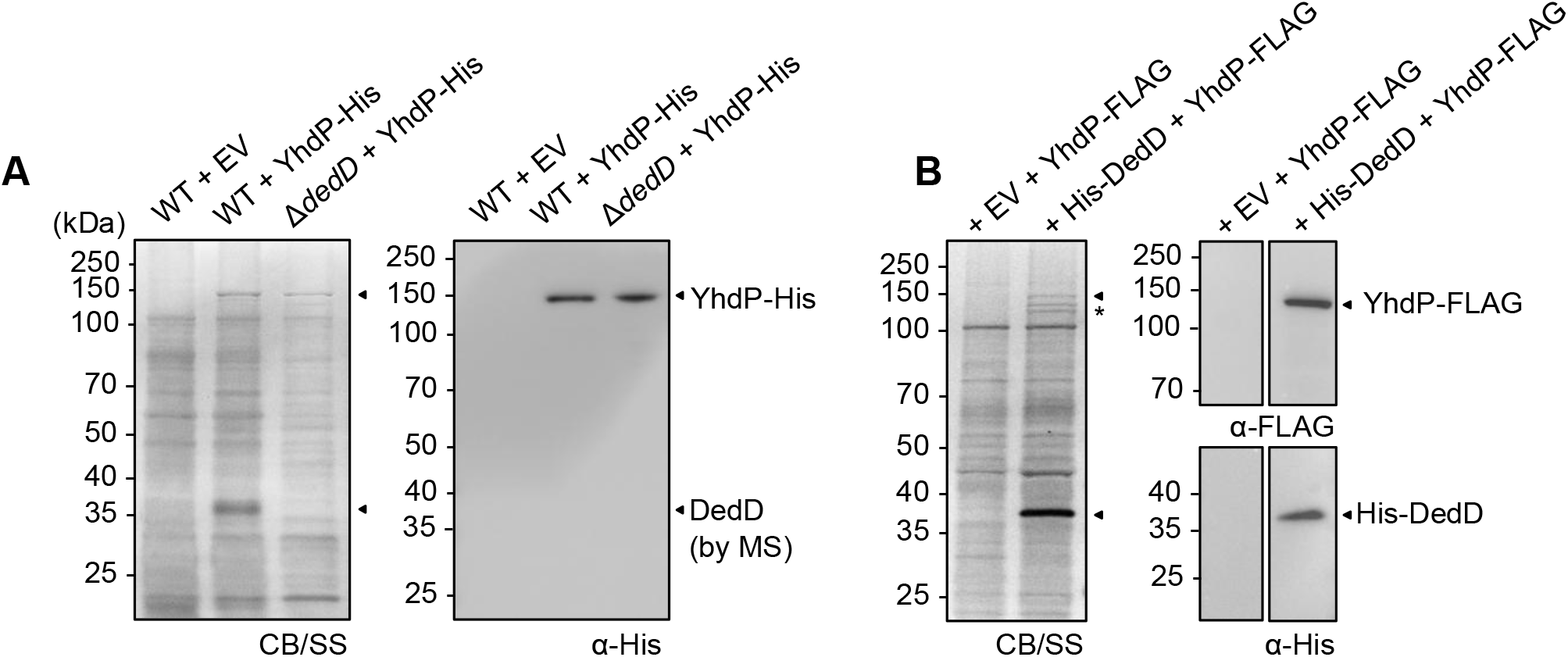
YhdP interacts specifically with DedD. (A) Affinity purification of WT or Δ*dedD* strain harboring vector or pET23/42-*yhdP*-*His*. MS/MS sequencing of the indicated band revealed DedD as the interacting partner. (B) Reciprocal affinity purification using strains harboring pBAD33-*His*-*dedD* (bait) and pET23/42-*yhdP*-*FLAG* (prey). Samples were subjected to SDS-PAGE stained with coomassie blue (CB) and silver stain (SS), and immunoblot analyses using α-His and α-FLAG antibodies. (*) Additional protein bands co-purified may represent native YhdP and/or C-terminal degradation resulting in loss of FLAG tag.

### YhdP localizes to the cell poles

DedD is a single-pass IM protein containing a periplasmic sporulation-related repeat (SPOR) domain (19), which is known to bind “denuded” peptidoglycan primarily at the cell septum; it localizes to mid-cell and influences the division process via an unclear mechanism (19). We therefore wondered if the observed interaction with DedD is important for the intracellular distribution of YhdP. To study localization, we constructed N-terminal sfGFP fusions of both DedD and YhdP, expressed from the pET23/42 vector. sfGFP-YhdP is fully functional in maintaining the OM barrier (Figure S3), and is capable of supporting growth of a Δ*yhdP* Δ*tamB* Δ*ydbH* strain (Figure S4). As expected, sfGFP-DedD retains its physiological localization to the cell septa during division (Figure 2A). In striking contrast, however, sfGFP-YhdP was found predominantly at the cell poles. We further demonstrated that this polar localization is not dependent on DedD (Figure 2A).

**Figure 2.**
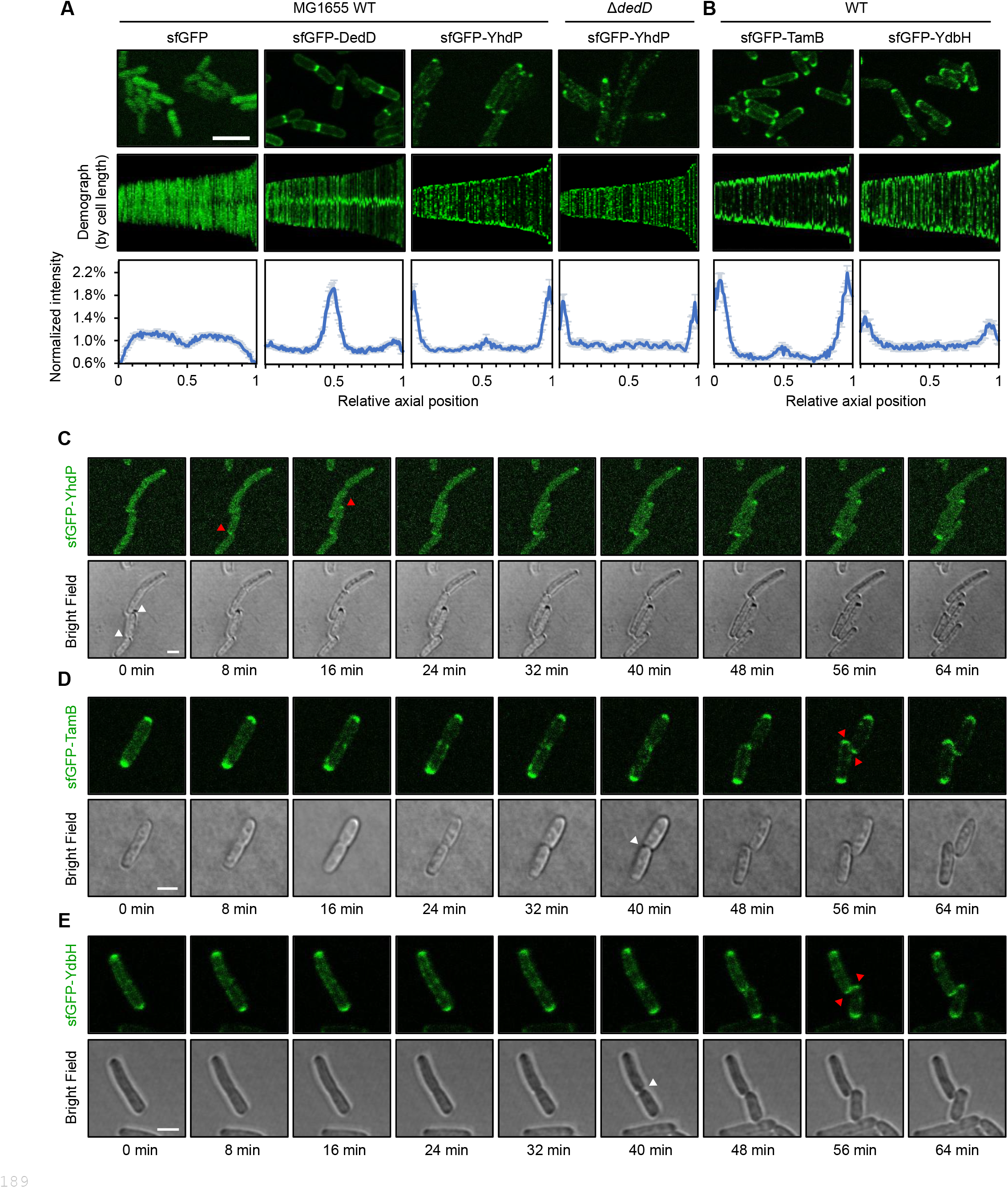
YhdP, TamB, and YdbH are enriched at the cell poles. (A),(B) (*top*) Fluorescence microscopy images of WT or Δ*dedD* cells harboring pET23/42-*sfgfp*, pET23/42-*sfgfp*-*dedD*, pET23/42-*sfgfp*-*yhdP*, pET23/42-*sfgfp*-*tamB*, or pET23/42-*sfgfp*-*ydbH*. Scale bar represents 5 µm. (*middle*) Demograph of the fluorescence sorted by cell length. (*bottom*) Distribution of the fluorescence along the long axis of cells (N=100). Error bars represent 95% confidence intervals. (C),(D),(E) Fluorescence time-lapse microscopy tracking the appearance of polar fluorescence signal in cell expressing sfGFP-YhdP, sfGFP-TamB, or sfGFP-YdbH. Images were originally taken at 2-minute intervals (see Supplemental Movies 1-3) and presented here at 8-minute intervals between frames. Scale bars represent 2 µm. White arrow(s) indicates completion of cell division and generation of new pole, only after which the fluorescence signals for all three putative PL transporters started to accumulate (red arrows).

Cells expressing sfGFP-YhdP were slightly elongated (Figure S5), indicative of division defects. This phenotype was similarly observed in strains expressing YhdP-His (Figure S6). We reasoned that expressing YhdP *in trans* might sequester DedD in these cells. Consistent with this idea, introducing additional copies of DedD partially reversed the elongated cell phenotype (Figure S5). While DedD and YhdP do not appear to co-localize, there are functional implications for their observed interaction.

This division defect suggests that the native ratio of YhdP to DedD was perturbed. We therefore constructed a strain where functional sfGFP-YhdP was instead produced from the original *yhdP* locus (Figure S7). When expressed at truly native levels (i.e. weaker signals), sfGFP-YhdP was still quantifiably spatially enriched at the cell poles (Figure S7). We conclude that polar localization of YhdP is relevant physiologically.

### TamB and YdbH also localizes predominantly to cell poles

The revelation that YhdP is found at the cell poles led us to investigate the subcellular localization of the other two putative PL transport proteins. We constructed N-terminally sfGFP-tagged versions of TamB and YdbH, and demonstrated their functionalities (Figure S3 and S4). Expressing sfGFP-TamB or sfGFP-YdbH did not result in division defects in WT cells (Figure S5). Interestingly, they both appeared strongly enriched at the cell poles (Figure 2B). We also observed localization of TamB at the mid-cell in longer cells, though it was difficult to differentiate whether the protein appeared there before or after complete cell separation (Figure 2B). Similar to sfGFP-YhdP, sfGFP-TamB remained polarly localized when it supported growth as the only functional PL transporter in cells (i.e. in Δ*yhdP* Δ*tamB* Δ*ydbH* background) (Figure S4). The strain expressing only sfGFP-YdbH exhibited expected morphological and OM barrier defects (Figure S4), as previously reported for the Δ*yhdP* Δ*tamB* double mutant expressing native YdbH (5, 6). We noted that sfGFP-YdbH was no longer polarly enriched in this Δ*yhdP* Δ*tamB* Δ*ydbH* strain (Figure S4), alluding to an interesting correlation between localization and morphology in this case. Taken together, these observations established that polar localization is a characteristic feature of the three collectively essential AsmA-superfamily proteins in *E. coli*, implying that their possible role(s) in PL transport is likely spatially confined.

### The putative PL transporters are targeted directly to the poles

When and how YhdP, TamB, and YdbH are spatially enriched at the cell poles is unclear. These AsmA-superfamily proteins could be recruited directly to the poles. Alternatively, they may be first recruited to the mid-cell at the late stage of division, and then stay at the new pole. To ascertain if the polar signal(s) appears before or after the completion of cell division, we performed time-lapse fluorescence microscopy with otherwise WT strains. Interestingly, all three sfGFP-YhdP, sfGFP-TamB, and sfGFP-YdbH only accumulated strongly at the new poles after two daughter cells had clearly separated (Figure 2C-E, Supplemental Movies 1-3). Notably, these putative PL transporters contain a large periplasmic domain that spans the cell envelope, therefore are likely immobile once the protein bridges are formed, due to the presence of the peptidoglycan mesh in the periplasm. Consistent with minimal lateral diffusion, the fluorescence signals for all three proteins remained largely static once formed (Supplemental Movies 1-3). While the exact molecular bases for polar localization of the putative PL transporters remain to be investigated, we conclude that YhdP, TamB, and YdbH are directly targeted to the cell poles (instead of division sites) and remain immobile thereafter.

## Discussion

Despite the specific interaction, majority of YhdP and DedD do not appear to occupy the same physical subcellular space. DedD is also not required for YhdP function in maintaining the OM barrier (Figure S8). While DedD diffuses away from the division site as daughter cells split, a low, residual level of DedD is also present everywhere, including the poles (Figure S9, Supplemental Movie 4). We speculate that the polar subpopulation of DedD could interact stably with, and to support, YhdP, e.g. facilitating assembly through the cell wall mesh. The exact functional implication for YhdP-DedD interaction requires further investigation.

The observed polar localization of YhdP, TamB, and YdbH imply that (anterograde) PL transport to the OM may be spatially confined to the cell poles. Consistent with this idea, removing YhdP was shown to partially rescue the unique IM shrinkage phenotype that specifically occurs at the cell pole, presumably due to high PL flux, in a strain with OM lipid dyshomeostasis (20). Polarly-confined PL transport contrasts strikingly with the assembly of new LPS and OMPs, which occurs all around the cell except the poles (3, 4, 21). This spatial segregation of OM biogenesis pathways is intriguing. Unlike LPS and OMPs, PLs are much more diffusive, and can be effectively re-distributed in the inner leaflet of the OM from the poles to support OM expansion elsewhere during growth. Besides, having the major constituents of the two leaflets of the OM be assembled at distinct localities may be optimal for the establishment of OM lipid asymmetry; inserting PLs into the inner leaflet where the outer leaflet is already filled (with LPS) could minimize scrambling and prevent mislocalization of PLs. While the field continues to gather experimental evidence of AsmA-superfamily proteins moving PLs directly, it will also be critical to test and understand the evolutionary advantage(s) to localize PL transport to the poles.

## Methods

### Strains, plasmids, and growth conditions

*E. coli* MG1655 is used as the wild-type for all experiment. Deletion alleles were constructed using λ Red recombineering (22) and subsequently moved into desired background strain using P1 transduction. MG1655 *yhdP*::*sfgfp*-*yhdP* chromosomally encoded strain was constructed using an established negative selection protocol (23). Strains were grown in Luria-Bertani (LB) broth (1% tryptone and 0.5% yeast extract, 1% NaCl) at 37°C. Chloramphenicol (30 µg/ml) and/or ampicillin (100 µg/ml) were added when required to maintain plasmid(s). Strains and plasmids used are listed in Table S1 and S2 respectively. Plasmids were generated with standard Gibson assemblies of PCR product generated with primers listed in Table S3.

### Affinity purification and SDS-PAGE

For each strain, 1.5 L culture was grown in LB broth at 37°C to mid-log (no IPTG added in all cases, and 0.02 % arabinose added for strain with pBAD33-*His*-*dedD*). Cells were washed and lysed with three rounds of sonication on ice (38% power, 1 s pulse on, 1 s pulse off for 3 min) in the presence of 1 mM of PMSF, 100 µg/mL of lysozyme, and 50 µg/mL of DNase I. Membrane fraction was collected by centrifugation at 25000 x g for 1 h at at 4 °C, and solubilized with 20 mM Tris-HCl pH 8.0, 150 mM NaCl, 10 mM MgCl2, 10%(v/v) glycerol, 1%(w/v) n-dodecyl-β-D-maltoside (DDM, Calbiochem)) at 4 °C for 1 hr. The solubilized membrane fraction was loaded into a pre-equilibrated column packed with 1.5 mL of TALON cobalt resin (Clontech) and incubated for 1 h at 4 °C with rocking. The mixture was allowed to drain by gravity before washing with 10 × 10 mL of wash buffer (20 mM Tris-HCl pH 8.0, 150 mM NaCl, 10 mM MgCl2, 10%(v/v) glycerol, 0.05 %(w/v) DDM, 10 mM imidazole) and eluted with 4 mL of elution buffer (20 mM Tris-HCl pH 8.0, 150 mM NaCl, 10 mM MgCl2, 10%(v/v) glycerol, 0.05 %(w/v) DDM, 200 mM imidazole). The eluate was concentrated in an Amicon Ultra 10 kDa cut-off ultra-filtration device (Merck Millipore) by centrifugation to ~100 μL. Concentrate was mixed with equal amounts of 2× Laemmli reducing buffer, heated at 95 °C for 10 mins, and subjected to SDS-PAGE analyses and immunoblotting. YhdP and DedD are better stained by Coomassie Blue and silver stain, respectively, therefore the resulting gel was doubly stained with silver stain (SilverQuest™ Silver Staining Kit, ThermoFisher) followed by Coomassie Blue.

### Fluorescence and time-lapse microscopy

Cell morphology and fluorescence localization of proteins were captured using an Olympus FV3000 confocal microscope at 100x magnification. Early mid-log (OD~0.2-0.4) cells grown by subculturing overnight culture at 1:2000 dilution at 37°C (after ~7-8 doublings), were spotted on LB 1% agarose pad for image acquisition. Images were analyzed using ImageJ (FIJI) and the MicrobeJ plugin (24, 25) to generate the demograph and the average intensity at different relative axial position. For time-lapse, the same field of view was imaged at 2-minute intervals for 80-110 minutes. The resulting image sequences were corrected for bleaching using the “Histogram Matching Bleach Correction” function in ImageJ (FIJI).

### Immunoblot

SDS-PAGE gel was transferred onto polyvinylidene fluoride (PVDF) membranes (Immun-Blot® 0.2 μm, Bio-Rad) using the semi-dry electroblotting system (Trans-Blot® TurboTM Transfer System, Bio-Rad). Membranes were blocked using 1X casein blocking buffer (Sigma). Rabbit polyclonal α-GFP was acquired from Abcam. Luminata Forte Western HRP Substrate (Merck Milipore) was used to develop the membranes, and chemiluminescent signals were visualized by G:BOX Chemi XT 4 (Genesys version 1.3.4.0, Syngene).

### Efficiency of plating

Sensitivity towards bacitracin or vancomycin were tested by spotting serially diluted cultures of the respective strains onto agar plates with the different conditions. Overnight cultures were 10-fold serially diluted in 150 mM NaCl on 96-well plate. Five µl of each dilution were spotted onto the plates and incubated overnight at 37°C. All results shown are representative of at least three independent replicates.

## Acknowledgments

This work was supported by the Singapore MOH NMRC OF-IRG (MOH-000145) and the Singapore MOE ARF Tier 1 grant (NUS-FoS Preparatory Grant).

## References

1. H. Nikaido, Molecular basis of bacterial outer membrane permeability revisited. Microbiology and molecular biology reviews 67, 593–656 (2003).

2. W. B. Tan, S.-S. Chng, How bacteria establish and maintain outer membrane lipid asymmetry. Annual Review of Microbiology 78 (2024).

3. L. Dubois, A. Vettiger, J. A. Buss, T. G. Bernhardt, Using fluorescently labeled wheat germ agglutinin to track lipopolysaccharide transport to the outer membrane in Escherichia coli. mBio, e03950–03924 (2025).

4. S. Kumar et al., Immobile lipopolysaccharides and outer membrane proteins differentially segregate in growing Escherichia coli. Proceedings of the National Academy of Sciences 122, e2414725122 (2025).

5. M. V. Douglass, A. B. McLean, M. S. Trent, Absence of YhdP, TamB, and YdbH leads to defects in glycerophospholipid transport and cell morphology in Gram-negative bacteria. PLoS Genetics 18, e1010096 (2022).

6. N. Ruiz, R. M. Davis, S. Kumar, YhdP, TamB, and YdbH are redundant but essential for growth and lipid homeostasis of the Gram-negative outer membrane. MBio 12, e02714–02721 (2021).

7. M. Hanna, A. Guillén-Samander, P. De Camilli, RBG motif bridge-like lipid transport proteins: structure, functions, and open questions. Annual Review of Cell and Developmental Biology 39, 409–434 (2023).

8. S. Kumar, N. Ruiz, Bacterial AsmA-Like Proteins: Bridging the Gap in Intermembrane Phospholipid Transport. Contact (Thousand Oaks) 6, 25152564231185931 (2023).

9. B. F. Cooper et al., Phospholipid Transport Across the Bacterial Periplasm Through the Envelope-spanning Bridge YhdP. Journal of Molecular Biology 437, 168891 (2025).

10. Y. Lin, B. Corry, Simulations of TamB suggest it facilitates lipid transport from the bacterial inner membrane. bioRxiv 10.1101/2025.03.12.642747, 2025.2003.2012.642747 (2025).

11. Y. Kang et al., Structural basis of lipid transfer by a bridge-like lipid-transfer protein. Nature, 1–8 (2025).

12. S. Kumar, R. M. Davis, N. Ruiz, YdbH and YnbE form an intermembrane bridge to maintain lipid homeostasis in the outer membrane of Escherichia coli. Proceedings of the National Academy of Sciences 121, e2321512121 (2024).

13. J. Selkrig et al., Discovery of an archetypal protein transport system in bacterial outer membranes. Nature structural & molecular biology 19, 506–510 (2012).

14. X. Wang, S. B. Nyenhuis, H. D. Bernstein, The translocation assembly module (TAM) catalyzes the assembly of bacterial outer membrane proteins in vitro. Nature Communications 15, 7246 (2024).

15. T. Wu et al., Identification of a protein complex that assembles lipopolysaccharide in the outer membrane of Escherichia coli. Proceedings of the National Academy of Sciences 103, 11754–11759 (2006).

16. W. B. Tan, S. S. Chng, Primary role of the Tol-Pal complex in bacterial outer membrane lipid homeostasis. Nat Commun 16, 2293 (2025).

17. Z. S. Chong, W. F. Woo, S. S. Chng, Osmoporin OmpC forms a complex with MlaA to maintain outer membrane lipid asymmetry in E scherichia coli. Molecular microbiology 98, 1133–1146 (2015).

18. J. Abramson et al., Accurate structure prediction of biomolecular interactions with AlphaFold 3. Nature 630, 493–500 (2024).

19. S. R. Arends et al., Discovery and characterization of three new Escherichia coli septal ring proteins that contain a SPOR domain: DamX, DedD, and RlpA. Journal of bacteriology 192, 242–255 (2010).

20. J. Grimm et al., The inner membrane protein YhdP modulates the rate of anterograde phospholipid flow in Escherichia coli. Proceedings of the National Academy of Sciences 117, 26907–26914 (2020).

21. G. Mamou et al., Peptidoglycan maturation controls outer membrane protein assembly. Nature 606, 953–959 (2022).

22. K. A. Datsenko, B. L. Wanner, One-step inactivation of chromosomal genes in Escherichia coli K-12 using PCR products. Proceedings of the National Academy of Sciences 97, 6640–6645 (2000).

23. V. Khetrapal et al., A set of powerful negative selection systems for unmodified Enterobacteriaceae. Nucleic acids research 43, e83–e83 (2015).

24. J. Schindelin et al., Fiji: an open-source platform for biological-image analysis. Nature methods 9, 676–682 (2012).

25. A. Ducret, E. M. Quardokus, Y. V. Brun, MicrobeJ, a tool for high throughput bacterial cell detection and quantitative analysis. Nature microbiology 1, 1–7 (2016).

