## Supplementary Material for "Polar localization of putative phospholipid transporters in *Escherichia coli*"

**Affiliations:**

[#] These authors contributed equally

* To whom correspondence should be addressed.

**ORCID ID:**

**This PDF file includes:**

Figure S1 to S9
Table S1 to S3

**As separate files:**

Supplemental Movie 1 to 4

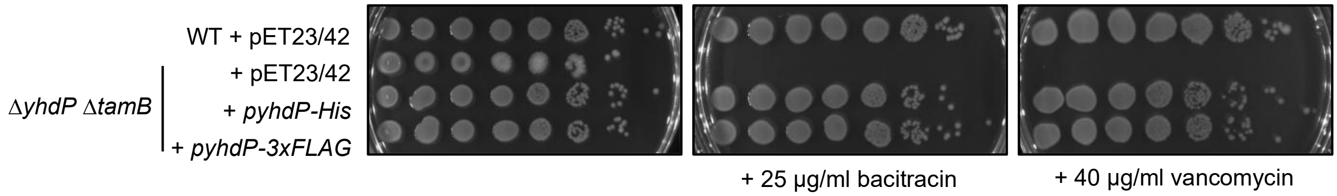


**Figure S1. YhdP-His and YhdP-Flag are functional.** Efficiency of plating (EOP) of MG1655 WT and Δ*yhdP* Δ*tamB* strains, expressing either YhdP-His or YhdP-FLAG on LB agar plates supplemented with bacitracin (25 µg/ml) or vancomycin (40 µg/ml) at 37°C. Cells with defective outer membrane (OM) barrier function become permeable to vancomycin and bacitracin, drastically sensitizing them towards these molecules. Δ*yhdP* Δ*tamB* double deletions result in OM barrier defects and sensitivity towards bacitracin and vancomycin (ref. 1); functional construct of YhdP restores the resistance.

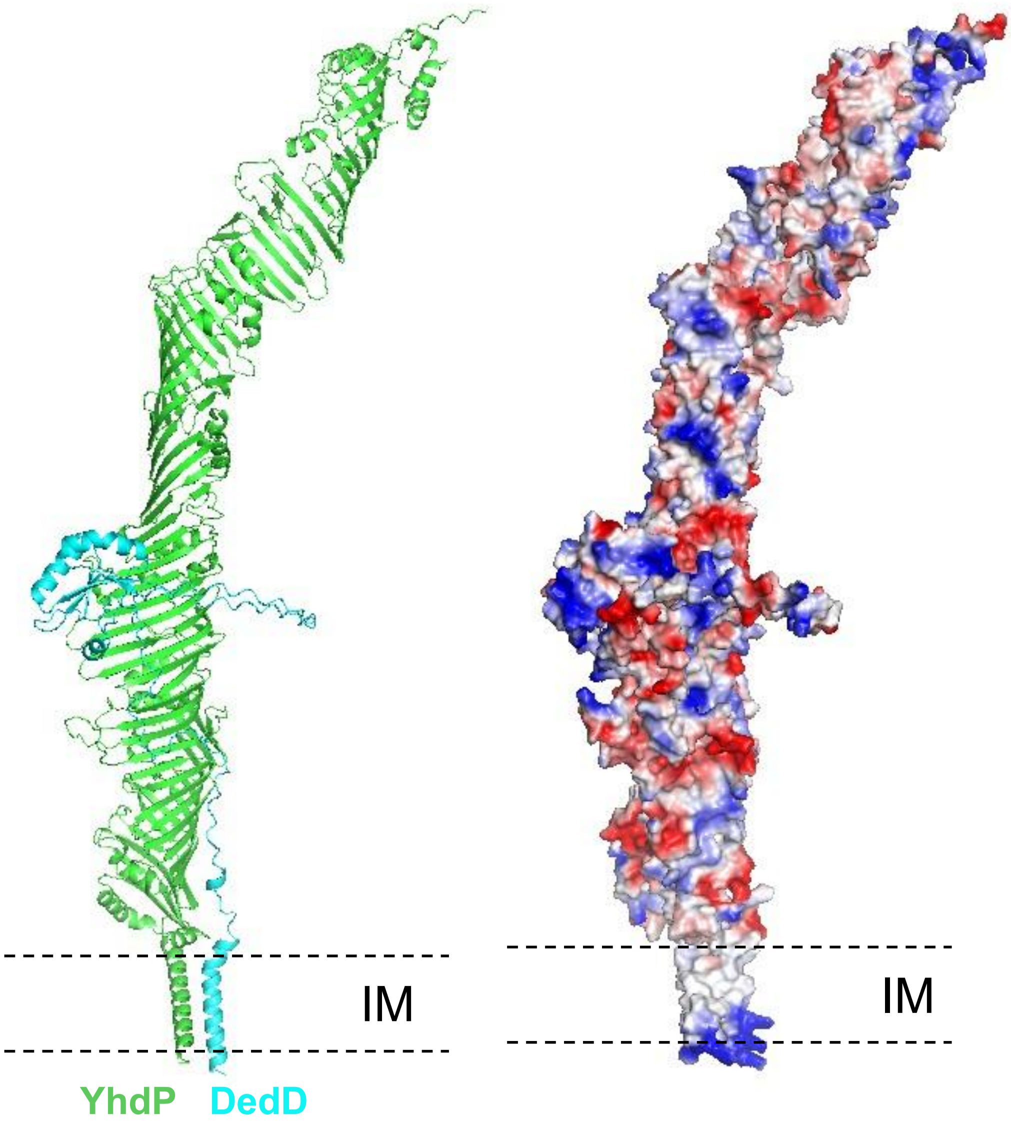


**Figure S2. AlphaFold3 prediction of YhdP-DedD heterodimer**. Possible structural model of a YhdP-DedD complex was predicted using the AlphaFold3 web server by inputting the YhdP and DedD protein sequences from MG1655. Resulting model of moderate confidence (ipTM = 0.6) suggests interaction interfaces at the transmembrane and periplasmic regions.

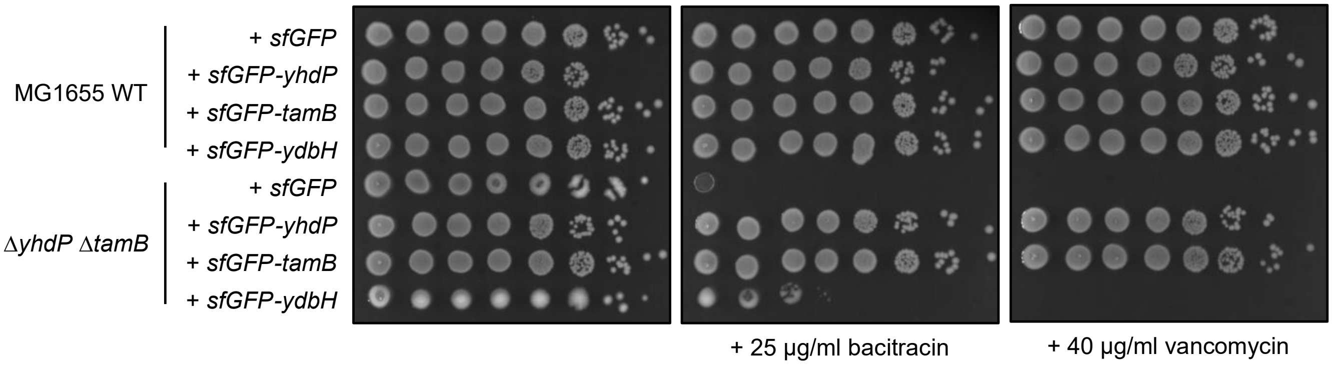


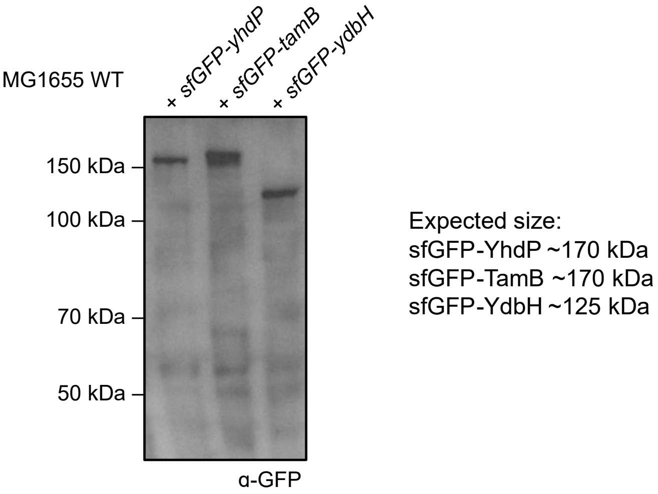


**Figure S3. sfGFP fusions of YhdP, TamB, and YdbH are functional.** (*Top*) EOP of MG1655 WT and Δ*yhdP* Δ*tamB* strains, expressing either sfGFP, sfGFP-YhdP, sfGFP-TamB, or sfGFP-YdbH, on LB agar plates supplemented with bacitracin (25 µg/ml) or vancomycin (40 µg/ml) at 37°C. Functional construct of YhdP or TamB restores resistance to bacitracin and vancomycin in the Δ*yhdP* Δ*tamB* double deletions strain at the concentration tested. Expression of functional YdbH is known to rescue outer membrane defects partially (ref. 3). (*Bottom)* Immunoblots of the whole cell lysate of the indicated strains using α-GFP antibody.

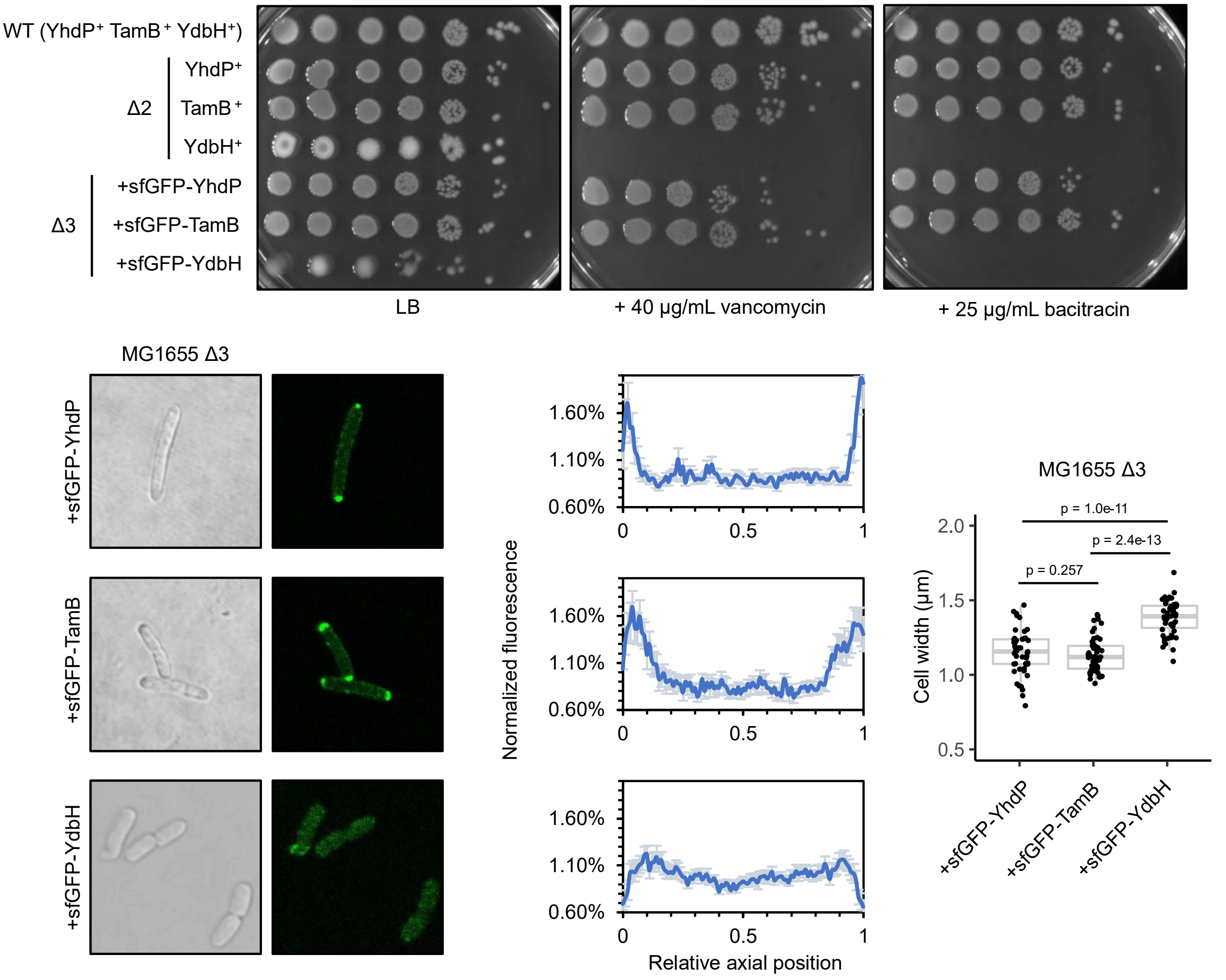


**Figure S4 Either one of sfGFP-YhdP, -TamB, or -YdbH is sufficient as the sole transporter to support growth.** (*Top*) EOP of MG1655 Δ3 (Δ*yhdP*::*frt* Δ*tamB*::*kan* Δ*ydbH*::*frt*) strains, expressing either sfGFP-YhdP, -TamB, or -YdbH from the respective pET23/42 plasmids, on LB agar plates supplemented with bacitracin (25 µg/ml) or vancomycin (40 µg/ml) at 37°C. The different plasmids were first transformed into an unmarked Δ*yhdP* Δ*ydbH* strain, where *tamB* was subsequently deleted using P1 transduction. Sensitivity to vancomycin and bacitracin of the Δ3 strains with either sfGFP fusion protein are identical to their corresponding Δ2 strain (with either native *yhdP*, *tamB*, or *ydbH* not deleted from the chromosome). (*Bottom*) Fluorescence microscopy images, distribution of the fluorescence along the long axis of cells (error bars represent 95% confidence intervals), and quantification of the cell width of the Δ3 strains (statistical significance tested with Wilcoxon-ranked-sum test).

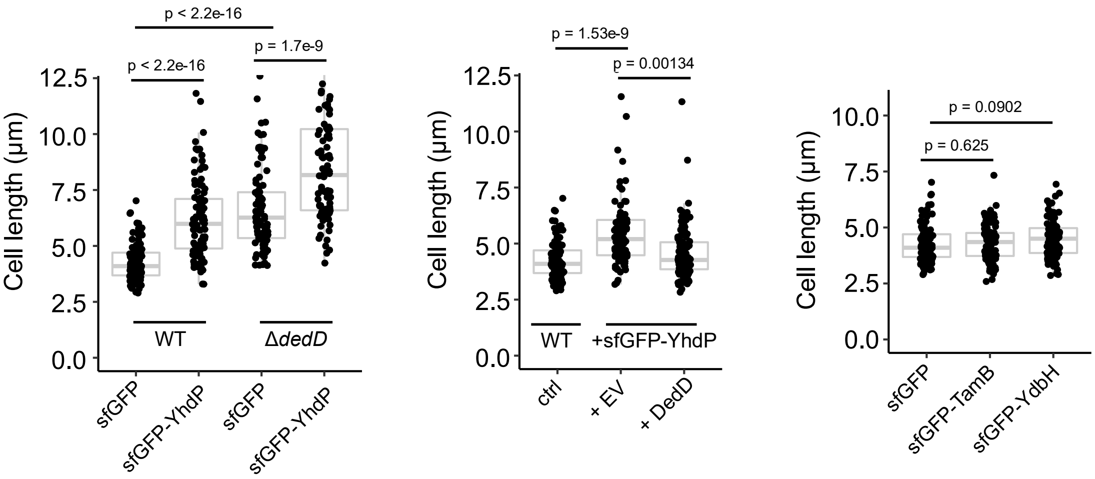


**Figure S5. Expression of sfGFP-YhdP causes mild division defects.** Cell length of the indicated strains was measured from the same samples used in Figure 2. Expression of sfGFP-YhdP (*left*), but not sfGFP-TamB or sfGFP-YdbH (*right*) resulted in longer cells, indicative of division defects. Expression of additional DedD reversed this morphological defect (*middle*). Statistical significance tested with Wilcoxon-ranked-sum test.

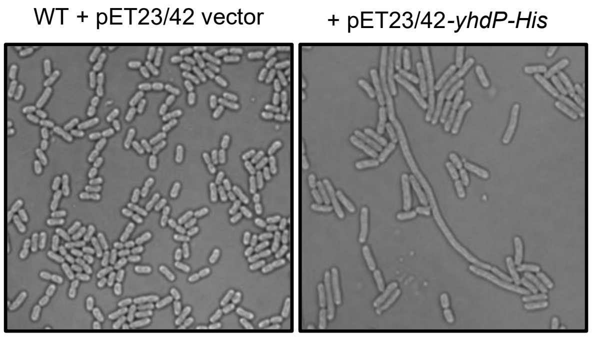


**Figure S6. Expression of YhdP-His also causes cell division defects.** Differential interference contrast microscopy images of WT strains containing either the pET23/42 empty vector or expressing YhdP-His from the same vector.

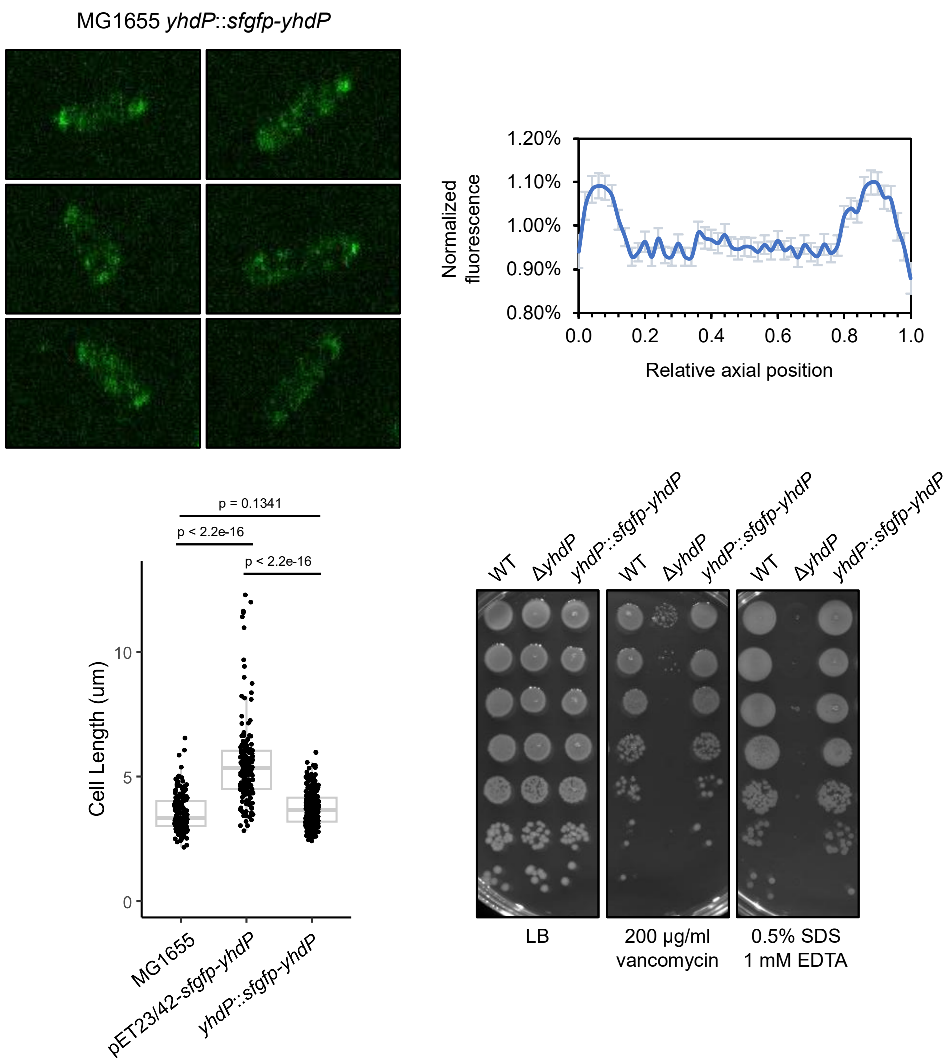


**Figure S7. Chromosomally-encoded sfGFP-YhdP exhibits polar localization.** (*top*) Fluorescence microscopy images, distribution of the fluorescence along the long axis of cells (N = 200, error bars represent 95% confidence intervals), and (*bottom left*) cell length quantification (statistical significance tested with Wilcoxon-ranked-sum test) of the chromosomally encoded *yhdP*::*sfgfp-yhdP* strain. (*bottom right*) EOP of MG1655 *yhdP*::*sfgfp-yhdP* strain in comparison with WT and Δ*yhdP* on LB agar plates supplemented with vancomycin (200 µg/ml) or 0.5% SDS 1 mM EDTA at 37°C.

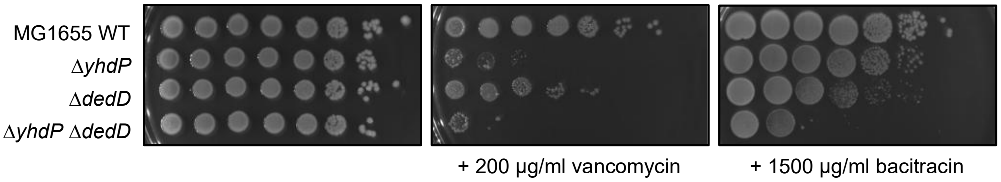


**Figure S8. Deletion of *dedD* does not result in OM defects similar to when YhdP is removed.** EOP of MG1655 WT, Δ*yhdP,* Δ*dedD,* and Δ*yhdP* Δ*dedD* strains on LB agar plates supplemented with bacitracin (1500 μg/ml) or vancomycin (200 μg/ml) at 37°C.

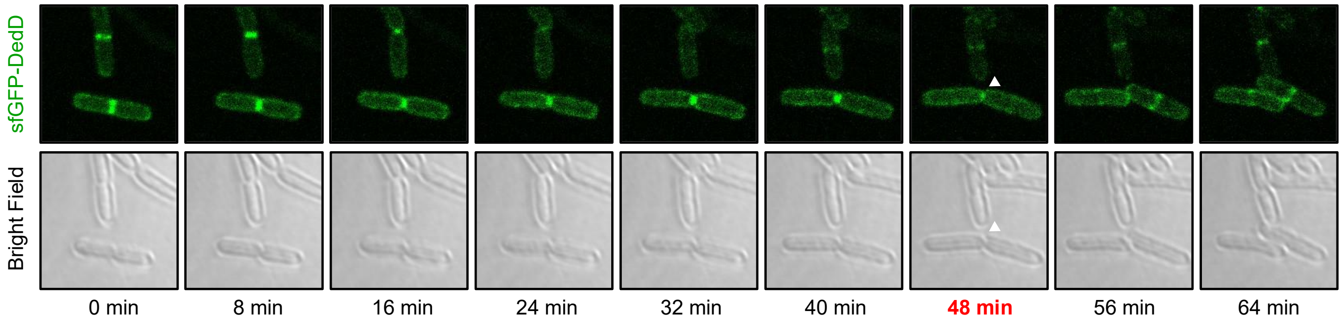


**Figure S9. Fluorescence time-lapse microscopy tracking the fluorescence signal of sfGFP-DedD.** Images were originally taken at 2-minute intervals (see Supplemental Movies 4) but presented here at 8-minute intervals between frames. White arrow(s) and red timepoint indicate completion of cell division and clear separation of daughter cells.

**Table S1. Strain list**

| Strain | Reference |
|---|---|
| MG1655 | WT |
| MG1655 Δ*dedD*::*kan*$^R$ | This study |
| MG1655 Δ*yhdP*::*frt* Δ*tamB*::*kan*$^R$ | This study |
| MG1655 Δ*yhdP*::*frt* Δ*ydbH*::*frt* | This study |
| MG1655 Δ*ydbH*::*frt* Δ*tamB*::*kan*$^R$ | This study |
| MG1655 Δ3 [Δ*yhdP*::*frt* Δ*ydbH*::*frt* Δ*tamB*::*kan*$^R$] * | This study |

***** MG1655 Δ3 genotype were constructed and maintained in the presence of pET23/42-*sfgfp-yhdP,* pET23/42-*sfgfp-tamB*, or pET23/42-*sfgfp-ydbH* plasmids.

**Table S2. Plasmid list**

| Strain | Resistance | Remarks |
|---|---|---|
| pET23/42 | Ampicillin | Empty vector (ref. 2) |
| pBAD33 | Chloramphenicol | Empty vector (ref. 3) |
| pET23/42-*yhdP-His* | Ampicillin | Pull down |
| pET23/42-*yhdP-3xFLAG* | Ampicillin | Reciprocal pull down |
| pET23/42-*sfgfp* | Ampicillin | Localization experiment |
| pET23/42-*sfgfp-dedD* | Ampicillin | Localization experiment |
| pET23/42-*sfgfp-yhdP* | Ampicillin | Localization experiment |
| pET23/42-*sfgfp-tamB* | Ampicillin | Localization experiment |
| pET23/42-*sfgfp-ydbH* | Ampicillin | Localization experiment |
| pBAD33-*His-dedD* | Chloramphenicol | Reciprocal pull down |
| pBAD33-3x*FLAG-dedD* | Chloramphenicol | Rescue of cell division defects |

**Table S3. Primers list**

| Primer name | Nucleotide sequence | Remarks |
|---|---|---|
| sfgfp-yhdP5 | gagctctacaaaGGATCCAGGCGATTGCCGGGGAT | Amplify *yhdP* from genomic DNA, with flanking homology to *sfgfp* (5') and pET23/42 backbone (3'), introduce two a.a. (glycine-serine between last a.a. of sfGFP and second a.a. of YhdP |
| sfgfp-yhdP3 | GTGGTGGTGCTCGAGtcaTTGCGCTTTTTCTTTACG | |
| pET2342sfgfp-yhdP5 | CGTAAAGAAAAAGCGCAAtgaCTCGAGCACCACCAC | Amplify pET23/42sfgfp backbone from pET23/42 (*sfgfp* inserted between NdeI and XhoI sites), with flanking homology to *yhdP* |
| pET2342yhdP3 | ATCCCCGGCAATCGCCTGGATCCtttgtagagctc | |
| sfgfp-tamB5 | gagctctacaaaGGATCCAGTTTATGGAAAAAAATC | Amplify *tamB* from genomic DNA, with flanking homology to *sfgfp* (5') and pET23/42 backbone (3'), introduce two a.a. (glycine-serine between last a.a. of sfGFP and second a.a. of TamB |
| sfgfp-tamB3 | GTGGTGGTGGTGCTCGAGctaAAACTCGAACTGATAGAGC | |
| pET2342sfgfp-tamB5 | GCTCTATCAGTTCGAGTTTtagCTCGAGCACCACCACCAC | Amplify pET23/42sfgfp backbone from pET23/42 (*sfgfp* inserted between NdeI and XhoI sites), with flanking homology to *tamB* |
| pET2342sfgfp-tamB3 | GATTTTTTTCCATAAACTGGATCCtttgtagagctc | |
| sfgfp-ydbH5 | gagctctacaaaGGATCCCTGGGTAAATATAAAGCCG | Amplify *ydbH* from genomic DNA, with flanking homology to *sfgfp* (5') and pET23/42 backbone (3'), introduce two a.a. (glycine-serine between last a.a. of sfGFP and second a.a. of YdbH |
| sfgfp-ydbH3 | GTGGTGGTGGTGCTCGAGtcaTTGTTTTTCCTCACACTC | |
| pET2342sfgfp-ydbH5 | GAGTGTGAGGAAAAACAAtgaCTCGAGCACCACCACCAC | Amplify pET23/42sfgfp backbone from pET23/42 (*sfgfp* inserted between NdeI and XhoI sites), with flanking homology to *ydbH* |
| pET2342sfgfp-ydbH3 | CGGCTTTATATTTACCCAGGGATCCtttgtagagctc | |
| sfgfp-dedD5 | gagctctacaaaGGATCCGCAAGTAAGTTTCAGAATCGG | Amplify *dedD* from genomic DNA, with flanking homology to sfgfp (5') and pET23/42 backbone (3'), introduce two a.a. (glycine-serine between last a.a. of sfGFP and second a.a. of DedD |
| sfgfp-dedD3 | GCCTAGGTATTAATCAAttaATTCGGCGTATAGCCC | |
| pET2342sfgfp-dedD5 | GGGCTATACGCCGAATtaaTTGATTAATACCTAGGC | Amplify pET23/42sfgfp backbone from pET23/42 (*sfgfp* inserted between NdeI and XhoI sites), with flanking homology to *dedD* |
| pET2342sfgfp-dedD3 | CCGATTCTGAAACTTACTTGCGGATCCtttgtagagctc | |
| yhdP::cat-tse2-5 | TTTTTGAGTCACATTTTTAGCAGACAAGGAGTGACGGgtgCTTCATTTAAATGGCGCGCC | Amplify *cat-tse2* (positive and negative selection marker) with flanking homology arms to yhdP locus, product transformed into MG1655 with lambda red helper plasmid for marking the *yhdP* locus |
| yhdP::cat-tse2-3 | GCGCGGCTCCAGTAAGCAGTAAAATCCCCGGCAATCGCCTCATATGAATATCCTCCTTAG | |
| H1-sfgfp-yhdP5 | GGGTTTTTGAGTCACATTTTTAGCAGACAAGGAGTGACGGatgtctaaaggtgaagaac | Amplify *sfgfp-yhdP* with homology arms to *yhdP* locus, product transformed into MG1655 with lambda red helper plasmid for marking the *yhdP* locus |
| yhdP3 | tcaTTGCGCTTTTTCTTTACGCGGTTGGCGC | |

**Supplemental Movie 1.** Fluorescence time-lapse microscopy video tracking the signal of sfGFP-YhdP at 2-minute intervals over 80 minutes. The movie displays each frame sequentially at the speed of 4 frames per second.

**Supplemental Movie 2.** Fluorescence time-lapse microscopy video tracking the signal of sfGFP-TamB at 2-minute intervals over 110 minutes. The movie displays each frame sequentially at the speed of 4 frames per second.

**Supplemental Movie 3.** Fluorescence time-lapse microscopy video tracking the signal of sfGFP-YdbH at 2-minute intervals over 110 minutes. The movie displays each frame sequentially at the speed of 4 frames per second.

**Supplemental Movie 4.** Fluorescence time-lapse microscopy video tracking the signal of sfGFP-DedD at 2-minute intervals over 110 minutes. The movie displays each frame sequentially at the speed of 4 frames per second.

**Supplemental references**


1. Ruiz, N., Davis, R.M. and Kumar, S., 2021. YhdP, TamB, and YdbH are redundant but essential for growth and lipid homeostasis of the Gram-negative outer membrane. *mBio*, *12*(6), pp.e02714-21.

2. Wu, T., McCandlish, A.C., Gronenberg, L.S., Chng, S.S., Silhavy, T.J. and Kahne, D., 2006. Identification of a protein complex that assembles lipopolysaccharide in the outer membrane of Escherichia coli. *Proceedings of the National Academy of Sciences*, 103(31), pp.11754-11759

3. Guzman, L.M., Belin, D., Carson, M.J. and Beckwith, J.O.N., 1995. Tight regulation, modulation, and high-level expression by vectors containing the arabinose pBAD promoter. *Journal of bacteriology*, *177*(14), pp.4121-4130.
